# Dynamic flux balance analysis of cancer metabolism using expression-based human cell line models

**DOI:** 10.64898/2026.10.15.759446

**Authors:** Liam P. Kelley, Ilija Dukovski, Darshi Shah, Louis Brezin, Kirill S. Korolev, Daniel Segrè

## Abstract

Aberrant metabolism that supports dysregulated growth is a key feature of cancer. Flux balance analysis, a computational approach that can predict genome-scale metabolic rates based on stoichiometry and mass conservation, has proven valuable in the prediction of pathway utilization and possible drug targets in cancer cells, but deep insight into growth dynamics under physiologically relevant conditions has been limited. To address this gap, we revisited the human metabolism model and its integration of gene-expression data, and we implemented simulations of metabolism in multiple cancer cell lines using static and dynamic flux balance analysis. Our models predict a substantial dependence of intracellular and exchange fluxes, and of the timing of resource utilization, on growth media and specific cell-line makeup. Secretions include lactate and other metabolites that could contribute to shaping the tumor microenvironment. Despite these differences in flux phenotypes, virtual genetic knockout screens predict high similarity of lethality across cell lines and media, and suggest a handful of possible cell-line specific targets. We finally demonstrate the power of dynamic simulations by setting virtual competition assays, which form the foundation for future extensions into dynamic simulations of tumor-stromal interactions in spatially-structured environment.

## Introduction

A primary hallmark of cancer is the dysregulation of cellular metabolism to support growth, proliferation, metastasis, and other key cancer phenotypes (1–6). This dysregulation includes unregulated uptake of glucose and amino acids, rewiring of central carbon metabolism to support biosynthesis, increased cycling of reductive and oxidative intermediates, and reshaping of the tumor microenvironment (7–9). A well known example of how cancer cells metabolically modify their microenvironment is the Warburg effect, where glucose is fermented to lactate despite sufficient oxygen for complete oxidation to CO_2_ (8, 10, 11). Despite decades of research, the mechanism behind and function of the Warburg effect are still not completely understood (10). The elevated consumption of glucose by cancer cells, however, has found applications in cancer diagnostics such as ^18^FDG-PET imaging (12, 13). In addition, various metabolic differences between healthy and tumor cells have been investigated as an avenue for inhibiting tumor growth or progression (14–18).

While studies of cancer metabolism have classically relied on *in vivo* and *in vitro* models, such as engineered mouse models and cancer cell lines, there is an increasing interest in the use of mathematical models and computer simulations as complementary tools to help understand cancer metabolism and prioritize possible drug targets (19–23). Whole-cell kinetic-based computational models of metabolism have been pursued for a few bacterial species (24). However, several challenges prevent the extension of this approach to human metabolism, including the lack of knowledge of kinetic parameters for most metabolic reactions (19, 22). The alternative approach of constraints-based modeling, often referred to as flux balance analysis (FBA), circumvents this limitation by imposing a steady state, and focusing on the prediction of reaction fluxes consistent with network stoichiometry and environmental boundary conditions (25, 26). In FBA, efficient optimization algorithms are generally used to identify specific cellular states associated with a given metabolic objective function (such as maximization of the growth rate). The stoichiometry for all reactions comprising the metabolic network of an organism (or cell line), together with a set of logical rules that define how specific genes map onto metabolic reactions, constitute the core knowledge encoded in a genome-scale metabolic model (GEM) (26). Despite the inherent complexity of mammalian metabolism, the idea that FBA could be leveraged to study human cells and tissues originated more than 20 years ago (27). Since the construction of the first genome-scale model for human metabolism (28), much progress has been made in creating increasingly complete and accurate human GEMs, with major releases including HMR1 (29), Recon2 (30), HMR2 (31), and most recently the HumanGEM models Human1 (32) and Human2 (33). These models have iteratively grown through the addition and revision of metabolic reactions and the inclusion of genetic regulation, ending up surpassing by almost an order of magnitude the complexity of bacterial GEMs (Fig. 1A). In the context of cancer biology, human GEMs have been used in multiple projects, including efforts to predict drug function and gene essentiality in hepatocellular carcinoma, renal carcinoma, prostate cancer, and others (34–36). Moreover, FBA-based computational studies sought to gain mechanistic insight into cancer metabolism, especially with respect to the role of the Warburg effect in invasion, growth, and ATP production (37–39).

**Fig. 1.**
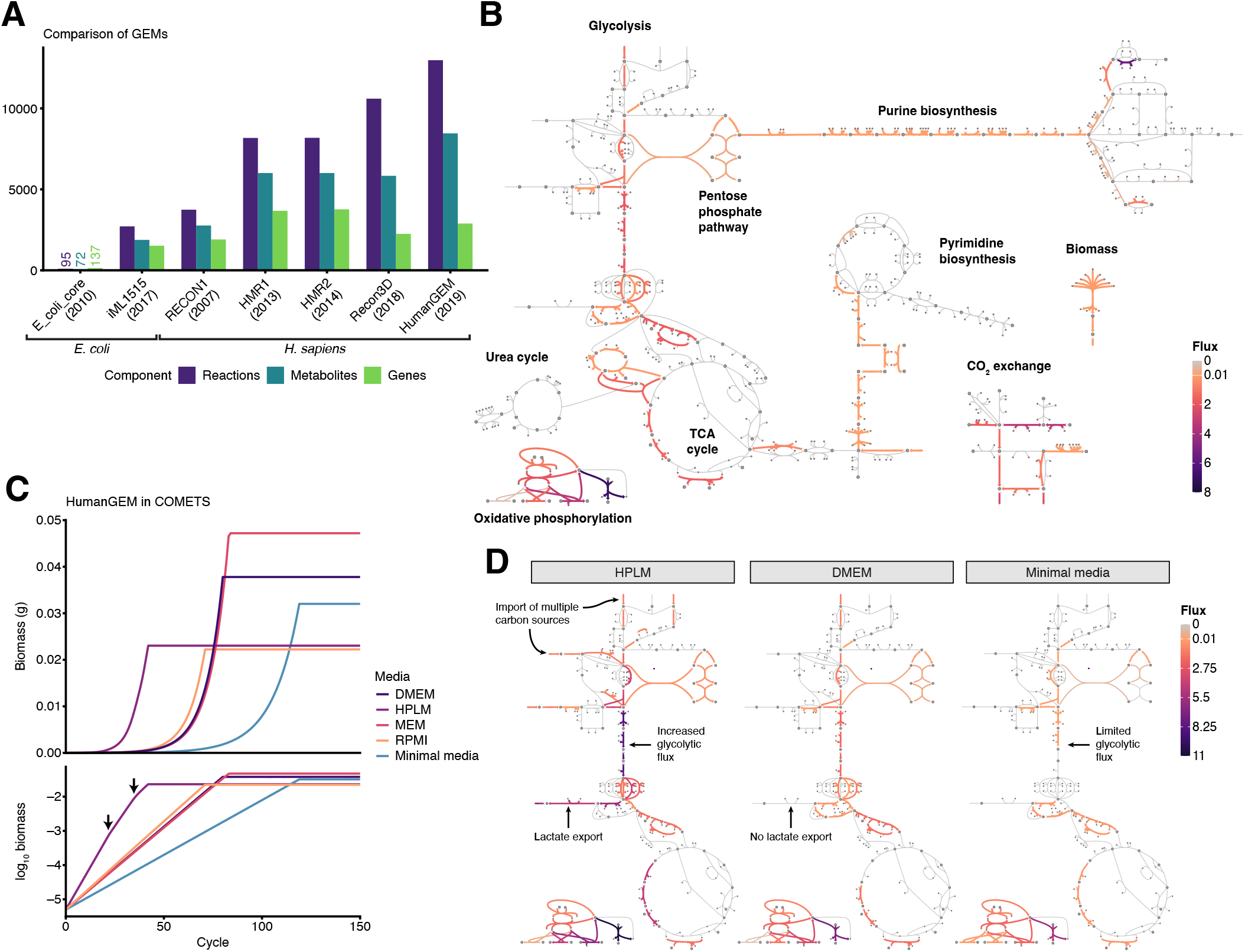
HumanGEM predicts differences in growth rate and carrying capacity between media based on internal fluxes. **(A)** Comparison of GEM complexity between two well-studied *E. coli* GEMs and major releases of human GEMs, including the Recon, HMR, and HumanGEM families. **(B)** Map of major pathways in HumanGEM, including central carbon metabolism, nucleotide biosynthesis, and electron transport. Overlaid on the map are fluxes from flux balance analysis of HumanGEM in DMEM media. **(C)** Biomass growth of HumanGEM in different media compositions as calculated by dynamic flux balance analysis within COMETS. **(D)** Escher metabolic flux maps for initial timepoints of dynamic flux balance analysis from (C), for the media conditions of HPLM, DMEM, and minimal media.

By extending FBA to sequences of intracellular steady states over many discrete time steps, dynamic FBA (dFBA) can generate predictions of how cellular biomass and environmental metabolites change over time (40, 41). In addition to being useful for predicting competition and cross feeding in microbial consortia (42, 43), dFBA has been coupled with biophysical models of biomass spread, metabolite diffusion, and chemotaxis, enabling simulations of multicellular dynamics in a spatially structured environment (44, 45). In particular, the COMETS platform (45, 46), initially developed for bacterial populations has been recently tested as a tool for studying human metabolism in health and disease, with viral infection as a first case study (47). However, the potential for dynamic flux balance modeling to help understand and forecast different types of cancer is still largely unexplored, with COMETS providing a valuable tool for computational investigations of human tumor metabolism in the context of spatially structured microenvironments.

In this paper, we sought to demonstrate the feasibility, versatility and possible applications of this methodology by implementing, within the COMETS platform, dynamic FBA simulations of human cancer tissues based on the HumanGEM reconstruction. Specifically, after testing and optimizing procedures for reproducible FBA and dFBA computation based on HumanGEM, we developed a new gene-expression-based protocol to construct cancer cell-line models, and generated extensive simulations of human cancer metabolism phenotypes. While results using the complete metabolic model failed to recapitulate several known cancer metabolism phenotypes, such as the Warburg effect, we observed extensive phenotypic diversity across cell line models and growth media, including production of lactate. Furthermore, dynamic FBA simulations in these cell line models highlight differences in growth rate and metabolite secretion, providing specific testable predictions amenable to future *in vitro* validation. Together, our results highlight the utility of context-specific models in detailed metabolic simulations of human cancers and serve as a crucial step in the development of accurate temporal multi-tissue simulations of cancer metabolism using dynamic FBA.

## Results

### Simulation of growth of HumanGEM in cell culture conditions

We began by working to simulate growth of the complete HumanGEM model on accurately recapitulated growth media. To systematically test and troubleshoot growth, we asked whether each biomass components was individually producible under each condition (see Methods). This test was performed in the presence of sets of extracellular metabolites representing common cell culture media, including Dulbecco’s modified Eagle medium (DMEM), minimal Eagle medium (MEM), RPMI 1600, human plasma-like medium (HPLM) (48), and an idealized complete medium containing all possible extracellular metabolites included in HumanGEM. This analysis highlighted a deficiency in the production of specific cofactors and lipids essential for biomass production under the existing media formulations (Fig. S1A), suggesting that some important media components were missing from the defined media compositions and were likely provided via serum, such as fetal bovine serum. A panel of extracellular cofactors were tested to identify the minimal set required for production of the entire cofactor pool (Fig. S1B). After identifying the non-essential fatty acid (NEFA) pool as capable of rescuing lipid pool production, individual components were tested as rescues, with the essential fatty acids linoleate and linolenate in combination rescuing production of the entire NEFA pool and therefore production of the lipid pool itself (Fig. S1C–D).

During this testing phase, we implemented further refinements to the HumanGEM model, based on newly identified issues, as reported in detail in a new dedicated GitHub repository, Human-GEM-custom. Main changes to the model included refinement of pathway for fatty acid metabolism, alongside edits to specific annotations to increase clarity and consistency (Fig. S1E). Escher maps (49) newly generated for this testing phase, ended up facilitating subsequent analyses as well. For purposes of simplicity, this model will be referred to as HumanGEM for the rest of this study.

Following the identification of missing media cofactors and essential nutrients, flux balance analysis was performed on HumanGEM to simulate growth on different media. As expected, major pathways in central carbon metabolism were highly activated when optimizing for biomass, with fluxes from glycolysis feeding into both the TCA cycle and the pentose phosphate pathway, alongside large scale fluxes through the electron transport chain and ATP synthase (Fig. 1B). We then performed dynamic flux balance analysis (dFBA) using the COMETS platform, and compared the growth of HumanGEM in different real-world media formulations. Dynamic FBA results displayed substantial differences in growth rate and total biomass yield when comparing standard media formulations alongside a minimal medium that consisted of only the essential nutrients required for biomass production (Fig. 1C). Mapping central carbon metabolism and electron transport chain activity highlighted distinct patterns of utilization of carbon catabolism between HPLM, DMEM, and minimal media, with import and partial oxidation of multiple carbon sources in the HPLM condition leading to export of lactate. In comparison, lactate export was not observed in the DMEM condition, and limited glycolytic activity was observed on the minimal medium (Fig. 1D). Our simulation results for the generic (i.e. not cell-type) version of HumanGEM suggest that human metabolism may display significant differences in pathway utilization as a function of medium composition.

### Generation of cell line models from transcriptional data

Following the initial characterization of HumanGEM using standard and dynamic FBA methods, we strove to generate context-specific models of HumanGEM. While HumanGEM is a valuable repository of the collective metabolic capabilities in human cells, additional constraints are required to accurately mimic any particular tissue, cell line, or developmental stage. For the purpose of this work, we specifically wanted to be able to compare simulated cancer metabolism across a few cell lines representative of specific tumors, grown in the different environment described above.

Multiple approaches were previously developed to generate tissue-specific human metabolism models based on different protocols for data integration (e.g., RIPTiDe (50) and ftINIT (51); see also comprehensive overview in Moyer *et al*. (52)). While some of these approaches were potentially consistent with our goals and with incorporation in COMETS, we designed a new method meant to be able to take advantage of large transcriptional datasets, and to incorporate human judgment in a final refinement step (see Methods and Discussion). Specifically, we relied on the Cancer Cell Line Encyclopedia (CCLE), which includes over 1700 cell lines analyzed using various approaches, including transcriptional profiling (53, 54). For subsequent work, and based on availability in this dataset, we focused on cancer cell lines A375 (melanoma), HeLa (uterine cancer), and MDA-MB-231 (breast cancer). We selected these cell lines as they capture a variety of tissue origins, growth rates, and metabolic phenotypes. In addition, they are commonly-utilized laboratory cell lines that will enable model tuning using *in vitro* experiments, and they are profiled using multiple omics techniques as part of CCLE. The workflow leverages expression data to classify genes in a binary on/off classification (similar to (51)), then utilizes gene-protein-reaction mapping in HumanGEM to identify reactions that would be missing in a specific expression pattern, and knocks those out of HumanGEM to create the cell line model (see Methods for details). When initially tested, cell line models produced from this workflow were unable to produce biomass, due to the knockout of essential genes. As a result, a select few necessary reactions were identified and re-introduced into the models to restore biomass growth (Fig. 2A). Following this steo of essential gene rescue, a total of 851 shared reactions had been removed from all models, suggesting that these reactions are connected to genes lowly expressed across different cell types (Fig. S2D).

**Fig. 2.**
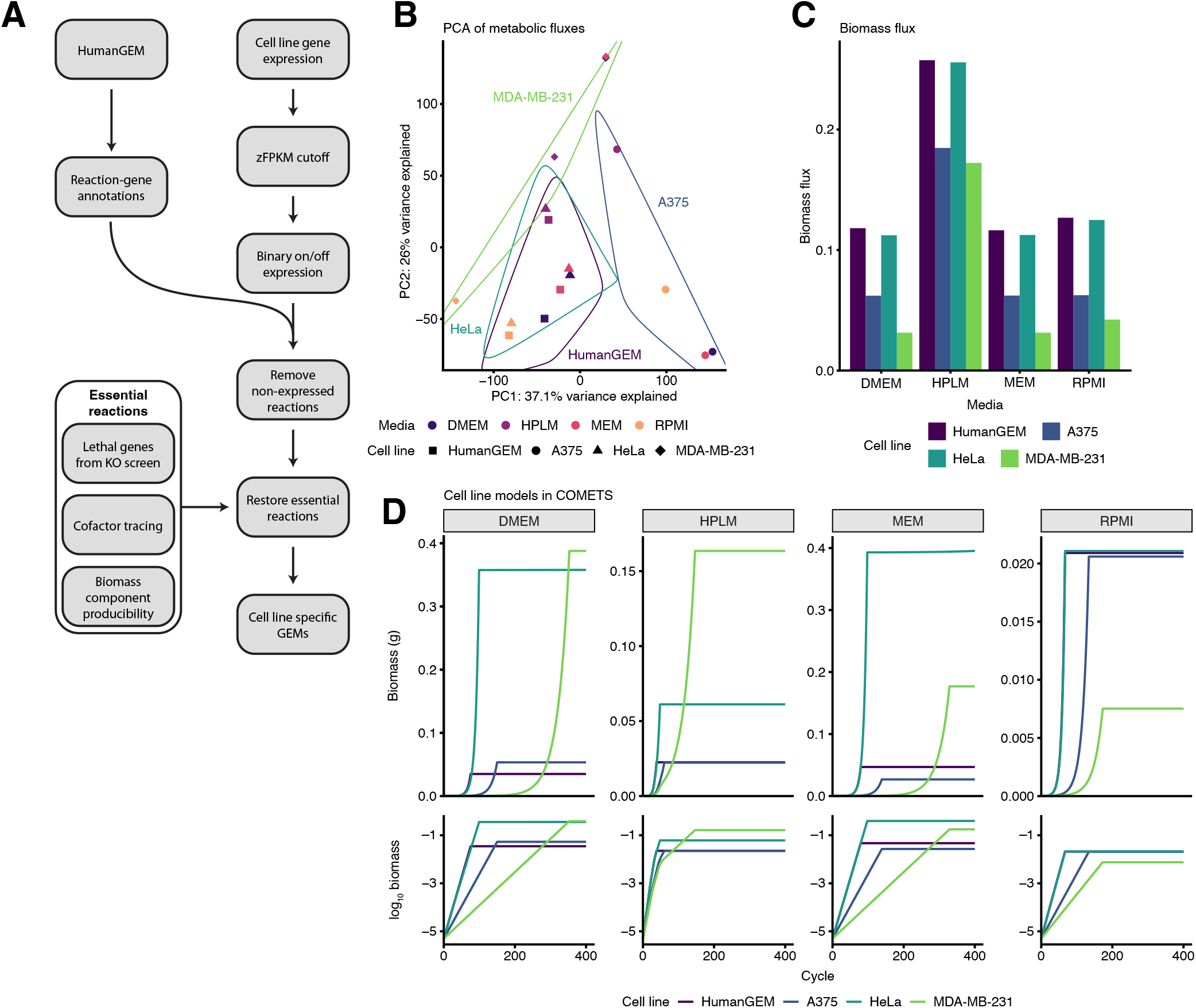
Cell line models generated from HumanGEM exhibit differences in both growth rate and total biomass yield. **(A)** Workflow of generation of cell line-specific models from HumanGEM using cell line gene expression from CCLE. **(B)** Principal component analysis (PCA) of cell line model fluxes generated by standard FBA in four media conditions. Points are colored by media condition and point shape denotes cell line model. Principal components 1 and 2 (PC1 and PC2, respectively) are plotted, with percent variance explained by each PC listed. **(C)** Biomass flux of cell line models in different media compositions as calculated by standard FBA. **(D)** Biomass growth charts for cell line models in different media compositions as calculated by dynamic FBA in COMETS.

With cell line models in hand, we then evaluated the metabolism of the models using both standard and dynamic FBA approaches in all four major media compositions. Principal component analysis (PCA) of all fluxes from standard FBA highlighted both media-dependent and cell line-dependent differences in metabolism in the dataset. All four cell lines, including the control HumanGEM model, were distinct by PCA when grown in the rich HPLM media. In contrast, the other three media conditions showed clustering largely driven by cell line, rather than media (Fig. 2B, Fig. S2E). The effect of HPLM was evident when plotting biomass flux in each cell line, with biomass flux significantly higher in HPLM across cell lines. Notably, the breast cancer cell line MDA-MB-231 exhibited much lower biomass flux in all non-HPLM media, suggesting a growth deficiency in that cell line relative to others in the panel (Fig. 2C). Notably, HeLa cells grew at nearly the same rate as HumanGEM, suggesting that they are lacking in very few metabolic pathways that contribute to maximal biomass flux. This is consistent with the finding that HeLa cells show limited differentiation and high stemness (55–57). Finally, we performed dynamic FBA of the cell lines using COMETS, based on standard parametrization of uptake kinetics (see methods), which may limit detailed quantitative insight into transient kinetics, but was shown to still generate insightful predictions (44–47). COMETS simulations highlighted stark differences in carrying capacity for the various cell lines in all media except RPMI. In both DMEM and HPLM, MDA-MB-231 cells had the highest biomass yield despite the slowest growth rate. In contrast, in both MEM and RPMI HeLa cells had the highest biomass yield. These data suggest that a tradeoff between growth rate and total biomass yield may be exhibited by cell lines in culture, with this tradeoff depending on nutrient availability in the media (Fig. 2D).

### Overflow metabolism of carbon byproducts and redox metabolites are altered in cell line models

Building on the differences in growth and biomass yield observed for the different cell line models compared to complete HumanGEM (Figure 2), we leveraged these cell line models to investigate additional metabolic characteristics associated with cancer metabolism phenotypes. In particular, we hypothesized that these different growth and yield properties would be reflected in different sets and rates of exudation and export of metabolites, including products of overflow metabolism. Among these, we were curious to explore differences in the production of lactate during aerobic growth, as a possible signature of the Warburg effect.

To generate a baseline of exudation against which to compare the different cell lines, we first examined the exports from control HumanGEM with FBA performed in DMEM media. In this case, we observed that major carbon-containing exports included carbon dioxide alongside carbonate, suggesting partial oxidation of carbon-containing substrates such as glucose were occurring. However, these were not the only outgoing flows: other secreted metabolites included acetaldehyde, a toxic oxidization intermediate responsible for hangovers (58); and 5-aminolevulinate (5-ALA) and porphobilinogen, heme pathway intermediates (Fig. 3A). In contrast, the principle export of MDA-MB-231 is lactate, consistent with partial oxidation of carbon-containing substrates as typified by the Warburg effect (Fig. 3B).

**Fig. 3.**
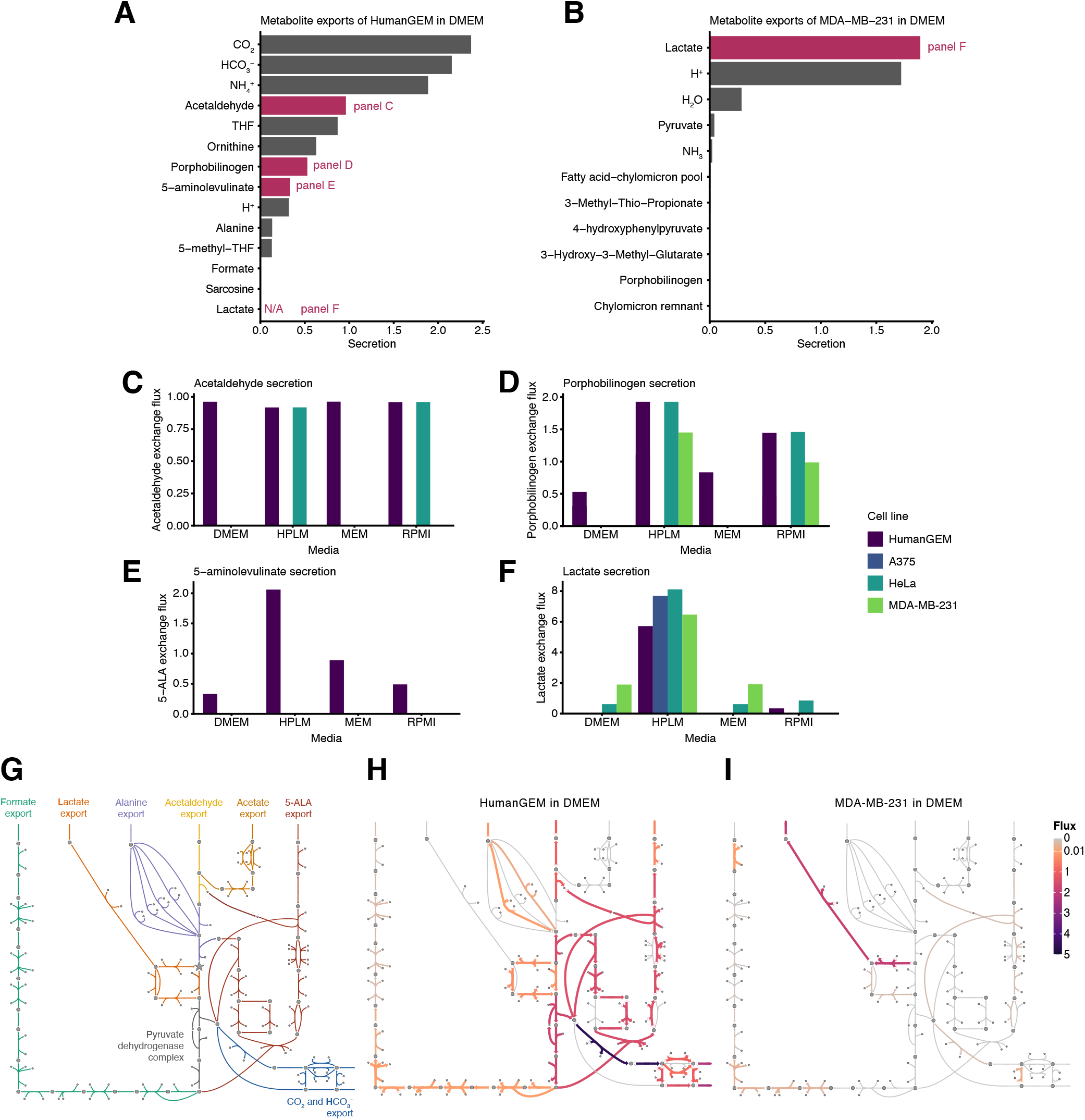
Prediction of overflow metabolism of carbon-containing compounds in HumanGEM is improved using cell-line models. **(A)** Metabolic exports of control HumanGEM when standard FBA is performed in DMEM media. Lactate has a net zero flux and is denoted as N/A for comparison to (B). **(B)** Metabolic exports of MDA-MB-231 cell line model when standard FBA is performed in DMEM media. **(C–F)** Exchange fluxes for (C) L-lactate, (D) porphobilinogen, (E) 5-aminolevulinate, and (F) acetaldehyde in HumanGEM and cell line models across media conditions. **(G)** Escher map of major pathways downstream of cytosolic pyruvate (denoted with a star in the center of the map) leading to export of carbon-containing compounds. **(H)** Map from (G) overlaid with fluxes from standard FBA of control HumanGEM in the DMEM media condition. Flux values are denoted via a color scale, with thin grey lines denoting zero flux. **(I)** Map from (G) overlaid with fluxes from standard FBA of MDA-MB-231 in the DMEM media condition.

Upon expanding the analysis to incorporate other cell lines and media conditions, we observed a shift in carbon handling between control HumanGEM and the cancer cell lines across media conditions. Notably, secretion of porphobilinogen, 5-aminolevulinate (5-ALA), and acetaldehyde were reduced in cell line models across media conditions. Acetaldehyde secretion was limited to HumanGEM in all media compositions and HeLa in HPLM and RPMI, with no secretion in either A375 or MDA-MB-231 (Fig. 3C). Porphobilinogen secretion was abrogated entirely in the A375 cell line (Fig. 3D). 5-ALA secretion was completely absent in all cell line models compared to control HumanGEM and across all media compositions (Fig. 3E). In addition, HPLM promoted the most lactate secretion, potentially due to the multiple carbon sources present in the media, including glucose, fructose, and glycerol (Fig. 3F). This phenotype was not observed in the original characterization of HPLM, however, suggesting that FBA’s prediction of import of multiple carbon sources may be inaccurate (48). Therefore, secretion of carbon-containing byproducts in the cell lines were more consistent with established cancer waste products than were the byproducts of the control HumanGEM model.

We visualized these differences in carbon metabolite exports using Escher. We began by generating a map tracing reactions leading from cytosolic pyruvate to the export of formate, lactate, alanine, acetaldehyde, acetate, 5-aminolevulinate, CO_2_, and HCO_3_ ^−^ (Fig. 3G). For HumanGEM grown in DMEM, fluxes from pyruvate predominantly led to CO_2_, HCO_3_ ^−^, and 5-aminolevulinate, as predicted from Fig. 3A (Fig. 3H). In contrast, cytosolic pyruvate was almost uniquely routed toward lactate production and export in MDA-MB-231 when grown in DMEM, highlighting the differences in carbon metabolism in the cell line model (Fig. 3I).

The substantial difference in lactate production between HumanGEM and MDA-MB-231 prompted us to ask whether specific reactions could be identified that markedly altered the fate of carbon exports derived from pyruvate. We therefore leveraged the map generated in Fig. 3G to identify down-stream reactions that could affect lactate export in the model. We identified 14 such reactions that could be required for export of one of the major carbon-containing compounds derived from pyruvate. Several reactions in these export pathways were missing in one or more of the cell line models relative to control HumanGEM, potentially explaining the observed differences in exports between HumanGEM and cell lines (Fig. S3). With the sole exception of pyruvate dehydrogenase complex (PDC), all reaction knockouts had no effect on biomass in control HumanGEM across media conditions, suggesting the reactions are completely dispensable for growth (Fig. S4A). While knockout of PDC reduced biomass flux in HumanGEM across most media, it also uniquely induced lactate secretion in DMEM and was also the knock-out with the highest lactate export in MEM (Fig. S4B). Expanding to look across cell lines, lactate export was unsurprisingly reduced or completely abrogated in the conditions targeting lactate (lactate exchange, cytosolic LDH, and mitochondrial LDH). Notably, MDA-MB-231 had high lactate export in all media except RPMI, highlighting media-dependent differences within cell lines. In general, cell lines demonstrated unique and media-dependent patterns of lactate secretion across the reaction knockout panel (Fig. S4C). Metabolic diversity of metabolic imports and exports was clear across the cell lines, with each cell line displaying a unique pattern of imports and exports in DMEM (Fig. S4D).

Another notable metabolite that was exported with an incongruously high flux by HumanGEM was tetrahydrofolate (THF), the reduced version of the necessary cofactor folate (Fig. 3A). We predicted that this large scale secretion of THF was a byproduct of high rates of folate reduction to balance cell redox state. Consistent with that hypothesis, we observed near-*V*_max_ import of folate in control HumanGEM and A375 in the DMEM and MEM media conditions. In contrast, this import was lower in both HPLM and RPMI media conditions and in the HeLa and MDA-MB-231 models (Fig. S5A). Notably, while both HumanGEM and A375 exhibited high rates of folate import, only HumanGEM exhibited commensurate rates of THF export (Fig. S5B). In HumanGEM, the vast majority of folate flux went into dihydrofolate followed by THF, with the produced THF exported from the cell rather than used to support biomass via the cofactor pool (Fig. S5C). We then investigated the metabolic dependency of HumanGEM on this folate uptake phenotype by limiting the upper bound of folate import, highlighting that THF was produced at a flux much higher than needed for biomass production (Fig. S5D). Given that reducing folate import had no impact on biomass production, we theorized that another redox reaction was being used to compensate. Indeed, we observed a compensatory increase in cystine import and cysteine secretion upon folate limitation, suggesting cystine was able to replace folate for the purpose of redox balancing (Fig. S5E–F). Together, the results of carbon-containing byproducts and redox intermediates highlight the refinement of predictions in over-flow metabolism enabled by the cell line models when compared to HumanGEM, and also suggest that multiple potential metabolic routes are available to cancer cells to achieve the metabolic requirements of carbon and redox balancing.

### HumanGEM and cell line models produce similar predictions in gene knockout screens

Based on the demonstrated metabolic differences between cell lines, we were next interested in whether these cell lines exhibited differences in gene essentiality that could be leveraged for potential treatments. To that end, we performed gene essentiality screens in control HumanGEM and the cell line models across media conditions, in a manner similar to previous approaches in human metabolic models (35, 59, 60).

After simulating knockouts of each gene in HumanGEM, we visualized the resulting fluxes using principal component analysis (PCA). PCA highlighted strong separation between the HPLM conditions and other media conditions in the panel, consistent with the results from standard cell line FBA in different media conditions (Fig. 4A, compare to Fig. 2B). Increased biomass flux and lactate secretion were evident in the HPLM conditions compared to other media conditions, again consistent with previous results (Fig. 4B–C).

**Fig. 4.**
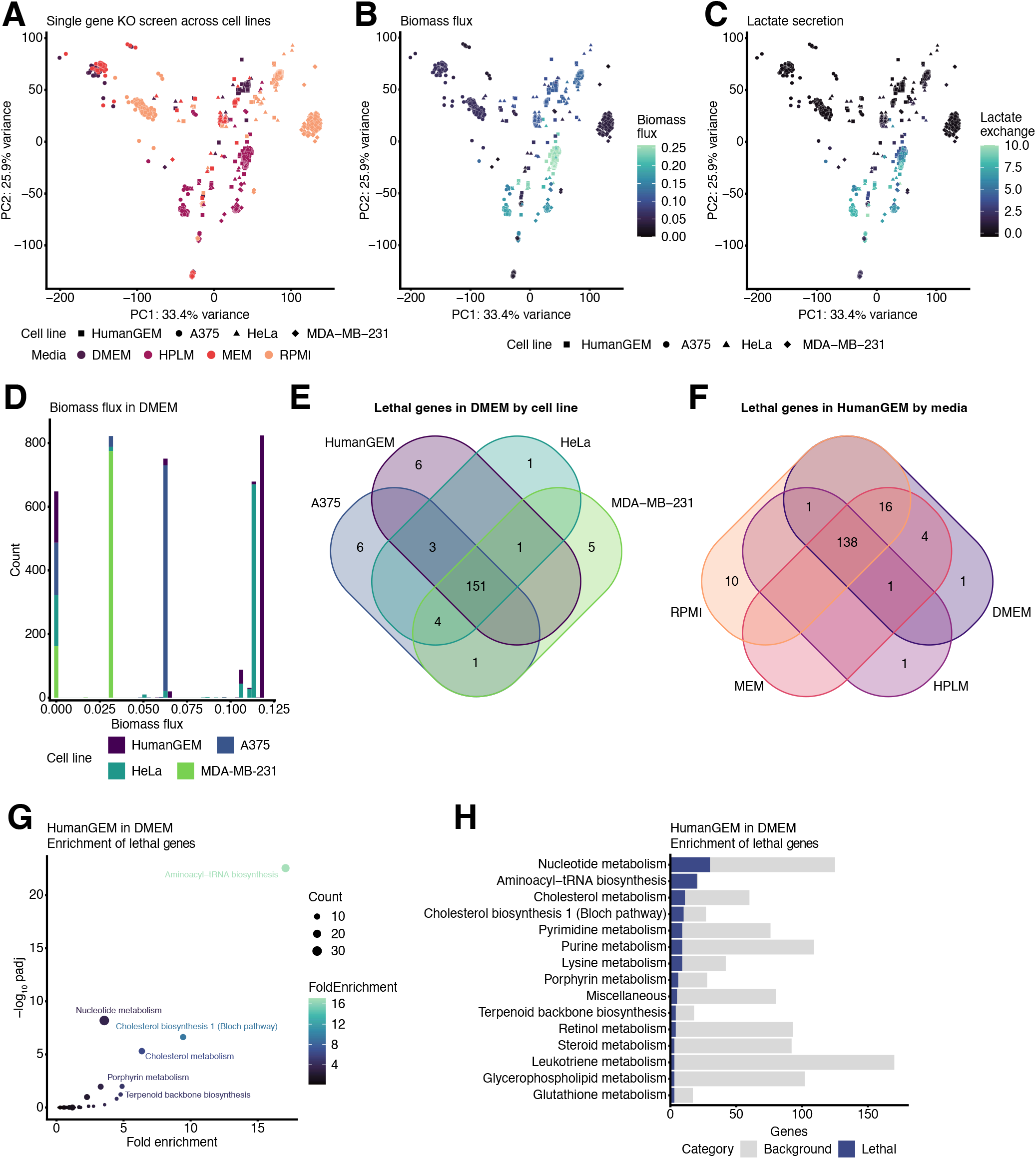
Gene knockout screens in HumanGEM and cell line models are largely consistent. **(A)** Principal component analysis (PCA) of single gene knockout screens performed in control HumanGEM and cell line models across media conditions. Points are colored by media condition, with cell line model denoted via point shape. **(B–C)** PCA from (A) with each point colored by (B) biomass flux rate or (C) lactate secretion rate, with cell line denoted by point shape. **(D)** Histogram of biomass flux rates across cell lines in the DMEM media condition. **(E)** Venn diagram of lethal genes (where knockout produces a biomass rate of zero) in DMEM by cell line. **(F)** Venn diagram of lethal genes (where knockout produces a biomass rate of zero) in control HumanGEM by media condition. **(G)** Pathway enrichment of lethal genes in control HumanGEM grown in DMEM condition. Fold enrichment is calculated as the ratio between the fraction of lethal genes in pathway and the total fraction of lethal genes. Pathways are plotted with adjusted *p* value (padj) and fold enrichment. **(H)** Top pathways from pathway enrichment in (G) by total number of lethal genes in pathway. Total number of genes in pathway is denoted by sum of blue and grey bars.

Given the pre-existing differences in biomass flux between cell lines (Fig. 2C), we then investigated the biomass flux distributions for each cell line in DMEM. Interestingly, the biomass flux distribution was largely penta-modal, with five distinct biomass flux values representing the control biomass flux for HumanGEM and each cell line, along with a biomass flux of 0 for all lethal knockouts. Notably, knockouts with reduced biomass fluxes for a given cell line often closely had similar biomass fluxes of a different cell line; for instance, knockouts in both A375 and HeLa produced biomass fluxes resembling control MDA-MB-231 (Fig. 4D). As a result, it appears as though specific reaction or pathway knockouts are largely responsible for the different growth rates of the cell line models, rather than large-scale differences in reaction availability in the models. While it has been demonstrated that loss of mitochondrial function in breast cancer inhibits growth, MDA-MB-231 does have evidence of functional electron transport, suggesting that the gene expression threshold for cell line generation may require adjustment (61).

As evidenced in the biomass flux histogram, a similar number of lethal genes were present in each of the four cell lines in DMEM. Comparing these lethal genes across models in DMEM, 151 lethal genes were shared across all models, with fewer than 10 lethal genes specific to a given model (Fig. 4E). Similarly, when looking at lethal genes in control HumanGEM across different media, 138 lethal genes were shared across all models, with a maximum of 16 shared across a subset of the media conditions (Fig. 4F). In general, biomass fluxes were strongly correlated between control HumanGEM and each of the cell line models in DMEM (Fig. S6A–C). Therefore, lethal genes were consistent across both cell line and media, suggesting that they arise from central pathways supporting biomass, rather than cell line- or media-specific discrepancies. To that end, we performed pathway enrichment analysis on lethal genes in control HumanGEM in the DMEM condition, which highlighted aminoacyl-tRNA biosynthesis and cholesterol metabolism as the most enriched pathways for lethal genes (Fig. 4G). By absolute number, meanwhile, nucleotide metabolism had the highest number of lethal genes (Fig. 4H).

While the majority of lethal genes were shared across cell lines, there were a limited number of genes that were lethal in one or more cancer model but not lethal in the control HumanGEM model. These cancer-specific targets represent potential therapeutic targets for cancer metabolism. As a result, we performed enrichment analysis of these cancer targets, finding a similar enrichment of cholesterol metabolism as in the general pool of lethal knockouts (Fig. S6D). Pathways with the most hits in this analysis also included retinol metabolism and vitamin E metabolism, suggesting specific differences to cofactor pathways in the cell line models alter lethality (Fig. S6E). As a result, few cancer-specific targets arose from the lethality screen.

In addition to characterizing lethal genes in the cell line models, we also characterized inhibitory genes, which are genes whose knockouts have nonzero biomass flux but reduce biomass flux by at least 1%. In contrast to lethal genes, there were fewer overlaps in inhibitory genes across cell lines and media conditions. Notably, few inhibitory genes were present in MDA-MB-231, resulting in the largest overlaps in inhibitory genes to be among control HumanGEM and HeLa, or HumanGEM, HeLa, and A375 together (Fig. S6F). However, strong overlap was still evident for inhibitory genes in HumanGEM across all four media compositions (Fig. S6G). Enrichment of these inhibitory genes showed clear enrichment of oxidative phosphorylation genes, with all ox-phos genes inhibitory in control HumanGEM when grown in DMEM (Fig. S6H–I).

Together, these findings reinforce that despite differences in reaction composition, both control HumanGEM and cell lines are still highly dependent on the synthesis of lipids, nucleotides, and proteins for biomass production under FBA rules. The similarity in gene essentiality identified in this approach could be a result of the limited number of cell lines used, or could arise from the dominant effect of biomass component biosynthesis in the base HumanGEM model that is revealed by FBA.

### Competition of cell lines in COMETS elucidates rules of metabolic competition in the human model

Beyond the differences in growth rate and metabolic fluxes observed across individual cell lines, one of the major questions dFBA simulations can help address is how genetically different populations of cells may interact in a shared environment. In the case of tumor modeling, these populations of cells may include tumor cells, stromal cells, and immune cells, among others. As a first step towards modeling metabolism in multiple interacting human cell types, we performed pairwise co-culture of our cell line models using COMETS. Specifically, we performed dFBA simulations starting with equal biomass amounts of a given pair of cell lines, and performed this analysis across all combinations of cell lines and media.

While COMETS can also simulate spatially structured environments, here we limited the analysis to non-spatial dFBA simulations.

We began by investigating the biomass amounts for each cell line at the end of the simulation, to determine whether the “winning” cell line would be determined by growth rate, final biomass yield, or other factors. When examining the fraction of total biomass for each cell line in pairwise competition, it became clear that the hierarchy of yield was HumanGEM, HeLa, A375, and then MDA-MB-231 (Fig. 5A). This order is the same as that of the overall growth rate of these cell lines in DMEM, as opposed to their total biomass yield when grown in isolation (Fig. 2C–D). When looking at how the fraction of each cell line changed over the course of the simulation, it was evident that the dominant cell line outcompeted the other cell line throughout the entirety of the simulation, rather than each cell line having a specific period of higher growth depending on nutrient availability (Fig. 5B). Thus, metabolic competition in this setting appeared to be an initial competition for major nutrients such as glucose and amino acids, with differential consumption of key nutrients playing little to no role in the outcome.

**Fig. 5.**
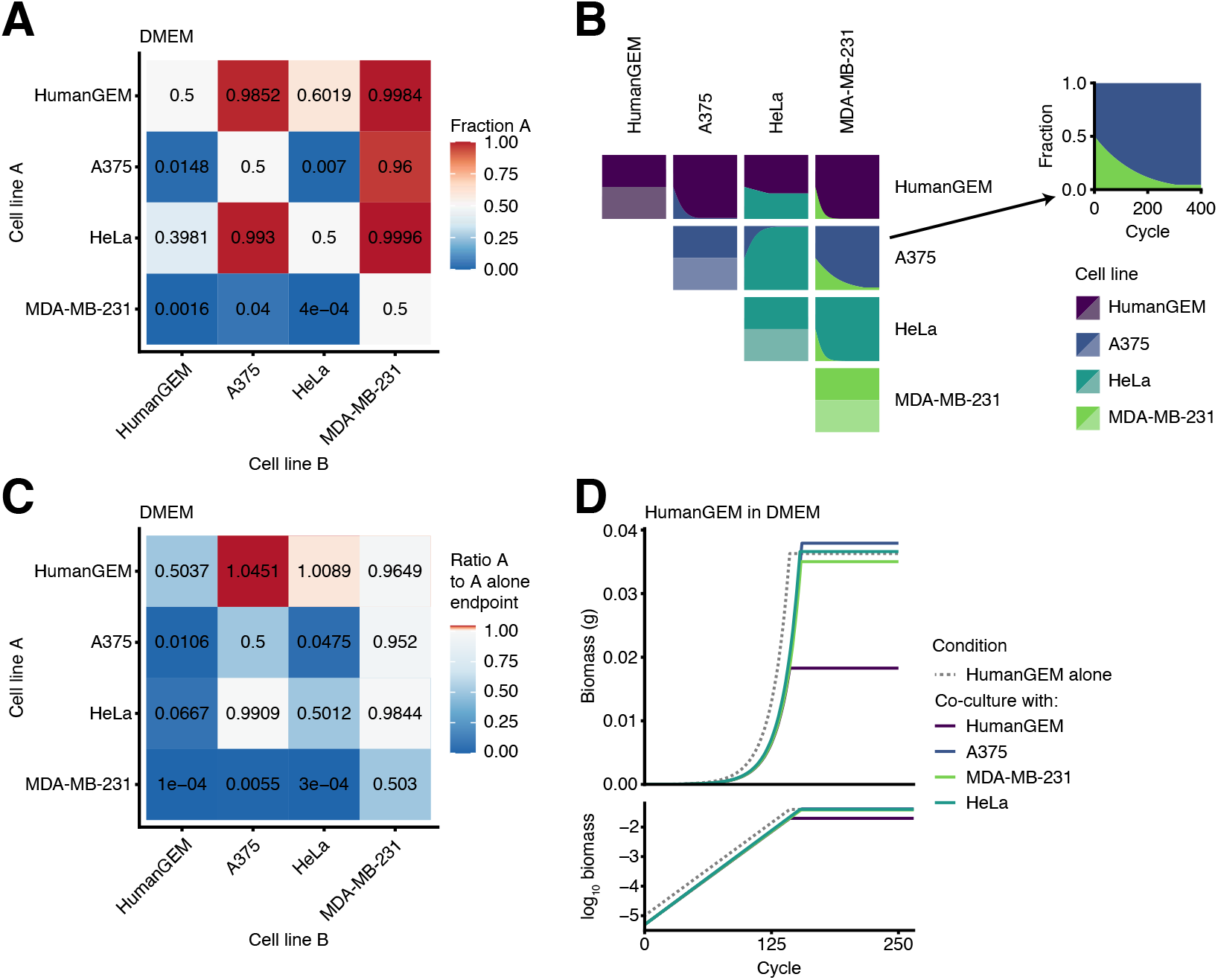
Initial growth rate determines outcome of cell line co-culture in COMETS. **(A)** Fraction of total well biomass at the final timepoint (cycle 200) for cell line A when grown with cell line B. **(B)** Plots of biomass fraction for pairwise co-culture of cell lines. Each square represents a line plot with *x* axis denoting cycle and *y* axis denoting biomass fraction. An example plot is shown on the right. **(C)** Ratio of endpoint biomass for cell line A when co-cultured with cell line B, compared to endpoint biomass of cell line A when grown alone. Blue colors denote growth below that of the cell line alone, while red colors denote growth above that of the cell line alone. **(D)** Biomass growth curves (both standard and log-transformed) for HumanGEM grow alone (dotted line) or co-cultured with the different cell lines.

Similarly, we investigated the effect of co-culture on the endpoint biomass of each cell line. To that end, we compared the endpoint biomass yield of each cell line grown alone to each pairwise co-culture combination. Surprisingly, control HumanGEM grew to a slightly higher endpoint biomass when co-cultured with A375 or HeLa than it did when grown alone (Fig. 5C–D), potentially suggesting a certain degree of cross-feeding. Other cell lines showed reduced growth when co-cultured compared to when grown alone, as expected. Therefore, while metabolic competition did not determine which cell line grew to a higher biomass yield, the yield itself was modulated by metabolic competition in a cell line-dependent manner.

## Discussion

In this study, we performed standard and dynamic flux balance analysis of a highly-detailed model of human metabolism, HumanGEM, along with cell line models generated from HumanGEM. We developed biologically-relevant simulations of human cancer metabolic dynamics that will enable validation of *in silico* predictions using *in vitro* experiments. We observed metabolic and growth differences between the models, including metabolic phenotypes consistent overflow metabolism and production of lactate. Together, these results form a key milestone toward dynamic simulations of multiple facets of cancer metabolism, and lay the groundwork for the development of more advanced context-specific metabolic models of tumors and surrounding tissue, enabling predictive simulations of the tumor microenvironment.

Genome-scale metabolic models aim to incorporate all genome-encoded metabolic reactions in an organism, generally lacking an explicit representation of regulation of these metabolic genes in specific cell types or during specific developmental stages of the organism (52). Context-specific subsets of GEMs, built through the integration of gene expression data, are key for studying cell- or tissue-specific metabolic phenotypes, and for modeling the diverse set of metabolic adaptations present in cancer (7–9, 62). Some prior efforts to generate context-specific subsets of human GEMs used enzyme expression levels to scale reaction limits, an approach that assumes high correlation between gene expression, protein abundance, and enzymatic flux despite evidence to the contrary (63, 64). To avoid this limitation, for our construction of cell line-specific models, we focused on a binary on/off classification of enzyme activity. As illustrated in in Table 1, our approach also differs from existing tools for analyzing GEMS in a context-specific manner, such as E-Flux (65, 66), scFBA (67), redHUMAN (68), RIPTiDe (50), INTEGRATE (69), and ftINIT (51). Notably, integration with COMETS for dynamic FBA requires the generation of standalone GEMs. Differences between these approaches are laid out in Table 1, highlighting our unique approach to context-specific GEM construction.

**Table 1.** Comparison of tools for context-specific GEM generation designed around transcriptional data and/or human models

| Tool | Input data | Role of expression data | Output data | Citation |
| --- | --- | --- | --- | --- |
| E-Flux | Transcriptomics (individual sample) | Set bounds | New constraints | (65) |
| E-Flux2 | Transcriptomics (individual sample) | Set bounds | Fluxes | (66) |
| scFBA | Transcriptomics (scRNAseq dataset) | Classify reactions, adjust reaction bounds | New constraints, smaller model | (67) |
| redHUMAN | Reaction thermodynamic data | N/A | Smaller model | (68) |
| RIPTiDe | Transcriptomics (single sample) | Weight fluxes | Smaller model | (50) |
| INTEGRATE | Transcriptomics, metabolomics, growth rates, measured fluxes | Adjust reaction bounds | New constraints | (69) |
| ftINIT | Transcriptomics or proteomics (individual sample or set of samples) | Classify reactions | Smaller model | (51) |
| This paper | Transcriptomics ($z$-scored from set of samples) | Classify reactions | Smaller model | |

Across standard and dynamic FBA, the generated cell line models showed marked differences in both growth and secreted metabolites compared to each other and to control HumanGEM. This is consistent with their differences in tissue of origin, as A375, HeLa, and MDA-MB-231 are derived from the skin, uterus, and breast respectively. Notably, the cell lines also highlighted differences in over-flow metabolism that is used to maximize growth in HumanGEM and HumanGEM-derived models. A principal difference arose with regard to carbon-containing exports. HumanGEM exhibited secretion of the carbon-containing compounds porphobilinogen, 5-aminolevulinate (5-ALA) and acetaldehyde. Porphobilinogen and 5-ALA are intermediates of heme metabolism; 5-ALA is associated with tumor metabolism, however rather than being secreted by tumors, it accumulates in the mitochondria and in fact forms the basis of a phototherapy technique to target cancer cells directly (70–72). Acetaldehyde, meanwhile, is the primary metabolite responsible for alcohol hangovers (58), and is linked to cancer as a carcinogenic metabolite, in particular to cancers of the digestive tract, rather than a byproduct of cancer (73–75). As a result, these carbon-containing exports in control HumanGEM are likely biologically inaccurate, with these predictions improved in the cell line models. Notably, while control HumanGEM exhibited no secretion of lactate in DMEM, lactate was the metabolite with the highest export flux in MDA-MB-231 in the same conditions (Fig. 3B).

Another metabolic adaptation present in control HumanGEM highlighted the central role of redox balancing in cancer metabolism. In control HumanGEM, folate was consumed at near maximal uptake flux (*V*_max_, see methods), with the majority converted to THF that was then secreted from the cell, thereby balancing redox state in the cell. Folate limitation and inhibition is a known mechanism for limiting cancer growth, and formed the basis of the first ever chemotherapeutic agents. However, this is due to the role of folate in nucleotide biosynthesis, rather than balancing of cell redox state (76, 77). Upon folate limitation, this balancing was then achieved via cystine import and subsequent reduction to cysteine, a pathway that is known to play a vital role in redox balancing in some cancers (78, 79). Thus, once again overflow metabolism in the complete HumanGEM model produced inaccurate predictions based on real requirements for growing cancer cells. Similarly, this phenotype was largely ameliorated in the cell line models, with HeLa and MDA-MB-231 not showing this folate-THF phenotype in any media condition.

In some cell line conditions, particularly in MDA-MB-231, the principal carbon export was lactate. This is consistent with the Warburg effect, which states that carbon cells selectively undergo partial oxidation of glucose into lactate despite sufficient oxygen for complete oxidation (3–6). Indeed, MDA-MB-231 cells imported less oxygen than was available, consistent with this phenotype. This may be due to missing mitochondrial complex genes, which contradicts findings that mitochondrial function is largely intact in cancers exhibiting the Warburg effect (10, 80). Computational studies have shown that the Warburg effect is advantageous in conditions of nutrient deprivation or in conditions where maximal ATP or biomass generation rate is required (37, 38, 81). As a result, though lactate generation from MDA-MB-231 is consistent with the Warburg effect, more work is needed to fully understand how the Warburg effect emerges in FBA in models of cancer metabolism.

The generation of cell line models from HumanGEM serves not only to enable a survey of different metabolic states in human metabolism via FBA, but also enables direct validation of FBA predictions in cell culture. A key regulator of metabolic predictions from FBA is metabolite import, which in this study was done with consistent Michaelis-Menten dynamics across all media metabolites. However, we plan to use *in vitro* cell line experiments to determine import parameters, using cell line growth and media metabolomics following glucose and amino acid modulation. Of particular interest is the relative depletion of essential amino acids in the media, which plays a key role in the prediction of final biomass yield in the different cell lines (Fig. 2D). Thus, the generation of virtual cell line models will enable cross-validation with real world metabolic data, improving the metabolic predictions for future simulations of 2D and 3D tumor dynamics.

## Supporting information

Supplement Table 1: media formulations

Supplement Table 2: cell line reaction knockouts

Supplement Table 3: cell line fluxes

## ACKNOWLEDGEMENTS

L. P. K., I. D., and D. Segrè were supported by grant 1R21CA279630-01 from the Informatics Technology for Cancer Research of the National Cancer Institute, and by the Rafik B. Hariri Institute for Computing and Computational Science and Engineering at Boston University. L. B. and K. S. K. were supported by grant 1R01GM138530-01 from NIGMS. The authors would like to thank the members of the Segrè lab for their thoughtful insights and discussion during the formulation of this work.

## Methods

### Curation of Human-GEM-custom

Modifications made to the existing version of HumanGEM, v1.19, were largely directed by the scope of the work. In particular, changes were made when gaps were noticed in pathways of interest at that particular time. Thus, each “release” of Human-GEM-custom largely focused on changes to a specific pathway, such as mitochondrial nitrogen handling, maintenance reactions, or fatty acid metabolism. All changes to Human-GEM-custom are tracked via git, and the repository for Human-GEM-custom can be accessed at https://github.com/segrelab/human-gem-custom. Major issues in the model were also submitted to the primary HumanGEM repository GitHub.

### Flux balance analysis

Standard flux balance analysis was performed using COBRA package version 0.29.0, running on Python version 3.10.5 (82). Unless otherwise stated, standard FBA refers to parsimonious FBA, where the optimizer objective (usually biomass) is first calculated, followed by a secondary step that optimizes for a minimal absolute value of fluxes within the model, representative of efficient use of enzymes by the cell(83).

Metabolic uptake for FBA was determined using Michaelis-Menten kinetics, which calculates the maximal rate of metabolic uptake by the model for each metabolite in the dFBA media. The formula is as follows:

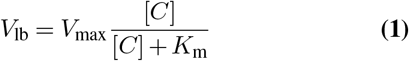

where *V*_lb_ represents the lower bound of the exchange reaction for a given metabolite based on concentration [*C*] and Michaelis constant *K*_m_. As a result, the exchange lower bound for each metabolite was set as *™V*_lb_, which for metabolites absent from the media corresponds to a lower bound of 0, preventing import. The upper bound was unchanged from model defaults, and was thus set at 1000.

### Optimization of linear programming accuracy

*V*_max_ for Michaelis-Menten uptake of media metabolites is based on biological principles of enzymatic activity, and relies on real-world data of enzymatic catalytic rate (such as *k*_cat_) for best accuracy. During FBA of HumanGEM, however, an issue was identified regarding the accuracy of linear programming tools used for FBA with various values of *V*_max_. In particular, *V*_max_ values that were too low, such as 10^*−*3^, led to solver inaccuracies, including loss of metabolite steady-state (metabolite net fluxes becoming non-zero) and out-of-bounds fluxes, manifesting in such ways as import of metabolites that were not present in the media. A comparison of multiple solvers, including Gurobi, GLOP, and GLPK, was performed in both COBRA and COMETS. The results highlighted that a *V*_max_ of 1 led to the most accurate solver behavior, minimizing out-of-bounds fluxes and metabolites out of steady state. For that reason, *V*_max_ = 1 for all analyses presented in the paper, unless otherwise stated.

### Dynamic flux balance analysis in COMETS

Single-well simulations in COMETS were performed with *V*_max_ = 1 and *K*_m_ = 1.5 *×* 10^*−*5^. Simulations were run using COMETSpy, a Python frontend for COMETS developed in tandem with COMETS (45). Analysis of COMETS biomass, fluxes, and media concentrations were performed following the runs using R.

### Formulation of replete cell culture media for growth of HumanGEM using FBA

Cell media formulations were derived from standard media formulations as available for consumers and commonly used in laboratories, as shown below.

| Media | Formulation source |
| --- | --- |
| DMEM | Gibco 11965 |
| HPLM | Sigma A4899 |
| MEM | Gibco 11095 |
| RPMI | Sigma R8758 |

These media formulations were then used to calculate exchange bounds for FBA, as described above. From there, the maximal production rate of each component of the biomass reaction was calculated as follows.

1. For metabolite *m*, create sink reaction *s* that can consume *m*
2. Set *s* as FBA objective
3. Run standard (non-parsimonious) FBA to calculate maximal flux through *s*

Sub-components of the biomass reaction were investigated for producibility in the same manner. The primary missing components from the biomass reaction were the cofactor and lipid pools (cofactor_pool_biomass and lipid_pool_biomass, respectively).

The cofactor pool was investigated using a combination of direct testing of extracellular cofactor rescues (*e*.*g*. testing biotin [e] to rescue synthesis of biotin [c]) and reverse flux tracing from a cofactor of interest to an extracellular metabolite. In addition, a large number of retinol derivatives are present in the cofactor pool, with retinol alone able to restore producibility of all of these species.

For the lipid pool, a component of the biomass reaction, flux tracing of the pool proved difficult due to the high rate of interconversion between lipid species in the model. Thus, an alternate approach was adopted. HumanGEM encodes 1625 extracellular metabolites, and we predicted a single metabolite would be able to restore lipid pool production. To this end, we tested pairwise combinations of pools of metabolites. As 1625 = 25 *×* 65, we generated pools of 25 metabolites and 65 metabolites, where each metabolite is assigned to one pool in different combinations. This is akin to assigning each metabolite to one spot in a 25 *×* 65 grid, and generating two sets of pools using rows and columns, respectively.

Therefore, if a given metabolite was sufficient to rescue lipid pool production, each of its respective pools would rescue lipid pool production. Using this approach, 234 metabolites matched the criteria of having both pools rescue lipid pool production. These 234 metabolites were then individually tested for rescue of all components of the lipid pool, and 14 rescued all components. Within those 14, one metabolite was of particular interest, as it was a defined fatty acid pool (the NEFA pool). Each component of the NEFA pool was then tested for ability to synthesize all members of the NEFA pool, with the combination of linoleate and linolenate able to synthesize all members of the NEFA pool, and therefore the lipid pool (Fig. S1C–D).

A minimal set of required nutrients was therefore assembled for all media formulations, and is listed in Table 2.

**Table 2.** Essential nutrients for cofactor and lipid pools in media formulations

| Nutrient | ID | Role |
| --- | --- | --- |
| α-tocotrienol | MAM01330e | Vitamin E |
| Aquacob(III)alamin | MAM01361e | Vitamin B<sub>12</sub> |
| Biotin | MAM01401e | Vitamin B<sub>7</sub> |
| γ-tocopherol | MAM01935e | Vitamin E |
| Lipoic acid | MAM02394e | Cofactor for pyruvate dehydrogenase |
| Pyridoxal-phosphate / pyridoxine | MAM02817e | Vitamin B<sub>6</sub> |
| Retinol | MAM02834e | Vitamin A |
| Riboflavin | MAM02842e | Vitamin B<sub>2</sub> |
| THF / folic acid | MAM01830e | Vitamin B<sub>9</sub> |
| Thiamine | MAM02982e | Vitamin B<sub>1</sub> |
| Linoleic acid | MAM02387e | Essential fatty acid |
| Linolenic acid | MAM02389e | Essential fatty acid |

### Generation of cell line models in HumanGEM

Cell line models in HumanGEM were generated using a combination of gene expression-based cutoffs for enzyme activity and rescues of essential reactions for viability.

Gene expression was determined using the Cancer Cell Line Encyclopedia (CCLE) RNA sequencing dataset (54). Raw counts were downloaded from the DepMap portal and combined with gene lengths (as calculated using the USCS

Genome Browser) to calculate FPKM values for each sample and gene in the dataset. Gene expression was classified using the zFPKM package, with the standard cutoff value of -3 for expression (84).

Following determination of binary (present/absent) gene presence for each gene in a given cell line, gene presence was then mapped to reaction functionality using gene-protein-reaction (GPR) rules in HumanGEM. In essence, gene status (T / True for present, F / False for absent) was inserted into gene-reaction formulae. As a generic example, consider a reaction that can be catalyzed using either of two different enzymatic dimers. The reaction gene rule could be listed as

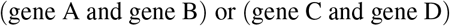

Replacing those with expression status (T or F) could produce, as an example

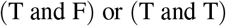

Using logic rules, this can then be simplified to True, meaning the reaction is functional and should remain in the model. This technique was applied to all genes in the model, and all reactions with gene rules, to determine whether the reaction remained active in the cell line. For reactions without gene rules, these reactions were assumed to be non-enzymatic and were kept active in the cell lines. This process produced a list of reactions to remove from the complete model in order to generate the cell line.

Essential reactions for model growth were reintroduced to the cell line models from multiple sources. The first source was from a reaction essentiality screen performed in HumanGEM in DMEM. All reactions that produced zero biomass when knocked out from HumanGEM were restored to the cell line models. Following rescue of those essential reactions, biomass producibility was evaluated using the same techniques as in Figure S1, highlighting that the cofactor and metabolite pools were often limiting in the cell line models. Metabolite tracing into production of heme, arginine, NAD^+^, vitamin D_3_, chromanol, and others was performed using fluxes from complete HumanGEM to identify necessary reactions to support synthesis of these cofactors and metabolites. After restoration of these essential genes, better viability was observed across the cell lines in the multiple media conditions. To improve cell line viability in the minimal media condition, a reaction essentiality screen was also performed in minimal media, with those essential reactions also restored in the cell line models. Finally, A375 cell line models were still deficient in vitamin D_3_ production, so manual testing identified MAR02138 as required to restore growth in that family of cell lines. Thus, a combination of reaction essentiality and manual tracing was done to rescue necessary reactions in the cell lines.

In essence, cell line models are represented as a list of reactions to knock out from complete HumanGEM, which is then done using COBRA or COMETS before running FBA using the model.

#### Generation of A375 cell line

The CCLE expression dataset included three distinct samples for the cell line A375, while the MDA-MB-231 and HeLa cell lines were represented with single samples. As a result, cell lines were generated for all A375 samples. A large number of reactions were knocked out in all cell lines, with 764 reactions shared across all cell line models (Fig. S2A). Expectedly, Jaccard analysis of the gene knockouts highlighted that the three A375 models were most similar to each other (Fig. S2B). After performing standard FBA in all media conditions, clustering analysis highlighted that the three A375 samples clustered with each other in all media conditions, suggesting that the three individual expression samples generated models with very similar metabolic behaviors (Fig. S2C). As a result, A375 sample #1 was selected as the representative model for A375, and was used for all further analysis.

### Gene knockout screens in HumanGEM

Gene knockout screens were performed using a similar approach to the generation of cell line models, as described above. For the complete HumanGEM model, gene knockouts were directly calculated using gene rules, as described above. For cell line models, gene knockouts were calculated by knocking out the gene of interest from the baseline gene expression used to generate the model. The new gene expression vector was then used to calculate reaction functionality as described above, using gene rules for each reaction. Any additional reaction knockouts compared to baseline were then stored for each gene knockout, and those reactions were removed from the baseline cell line model (after essential reaction rescue) to generate the knockout model for a given gene knockout. Thus, gene knockouts were calculated relative to the baseline expression for the cell line while still allowing for lethality in the knockout screen.

### Enrichment of knockout reactions

Enrichment of hits from the gene knockout screen in HumanGEM and cell lines was performed using clusterProfiler (85). Given enrichment was performed on reaction subsystems as defined within HumanGEM, while the knockout screen was done on genes, a mapping between genes and reaction subsystems was needed.

This mapping was generated by mapping genes to reactions, and then reactions to subsystems. For each gene, the subsystem mapping was generated as the union of all subsystems for that gene’s reactions. The majority of genes (1936 out of 2887) mapped to a single subsystem. Using this mapping, enrichment was then performed using the enricher function within clusterProfiler.

### Co-culture of cell lines in COMETS

For cell line co-culture competition assays in COMETS, a total of 10^*−*5^ g of biomass was placed in the COMETS layout “well”. Thus, for single cell line controls, the cell line had a starting biomass of 10^*−*5^ g, while for two cell line experiments, each had a starting biomass of 5 *×* 10^*−*6^ g. Simulations were run for 1000 cycles. Biomass outputs from COMETS were then imported into R for further analysis.

## Supplementary Figures

**Fig. S1.**
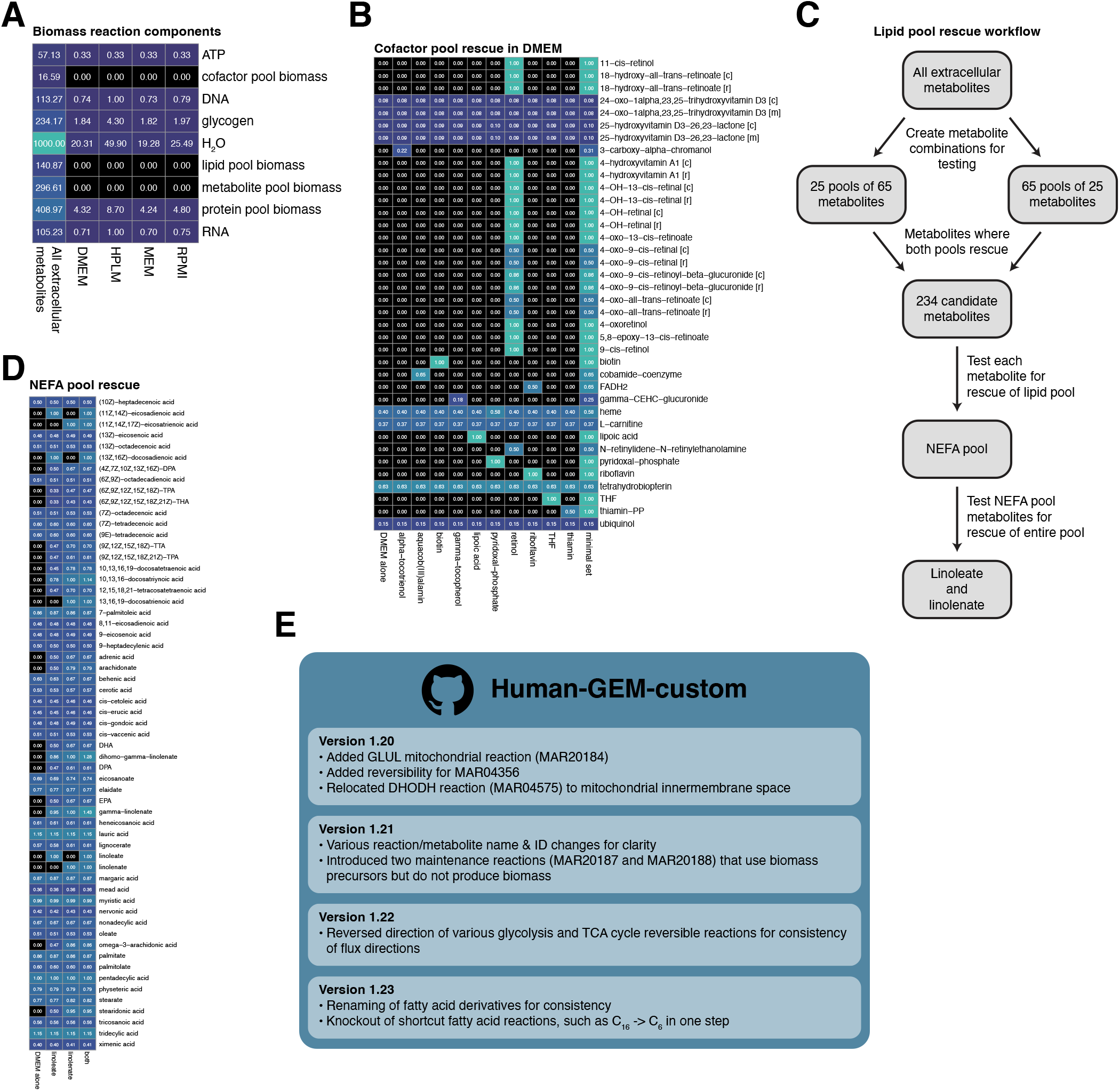
Identification of missing cofactors for growth of HumanGEM in media formulations. **(A)** Maximal rate of production of components of the biomass reaction under different media conditions, highlighting the infeasibility of production of the cofactor and lipid pools under initial media conditions. **(B)** Maximal rate of production of components of the cofactor pool in DMEM with addition of specific metabolites in the media. The minimal set is a combination of the other listed metabolites. **(C)** Workflow for the identification of the missing media components to rescue the feasibility of production of the lipid pool. **(D)** Maximal rate of production of components of the NEFA pool, highlighting that a combination of linoleate and linolenate were capable of rescuing all components of the pool. **(E) Major changes implemented in versions of Human-GEM-custom following the fork from v1.19**.

**Fig. S2.**
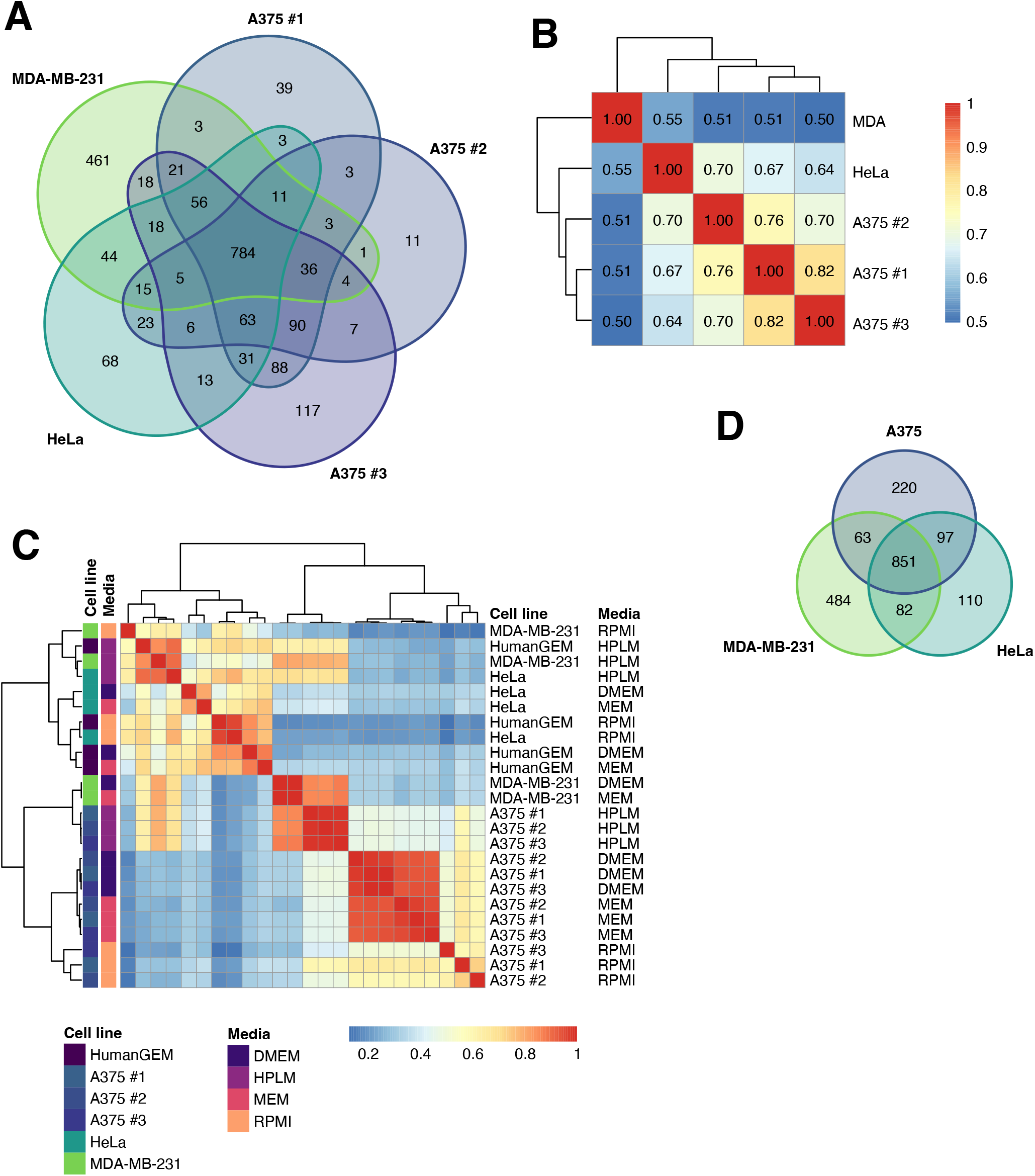
Evaluation of cell line specific models from HumanGEM. **(A)** Venn diagram of reactions removed from control HumanGEM to produce each cell line model (before rescue of necessary reactions). **(B)** Clustered Jaccard analysis of removed reactions from (A). **(C)** Clustered correlation heatmap of fluxes from standard FBA in each cell line model (and control HumanGEM) across media conditions. **(D)** Same Venn diagram as (A) after removal of A375 #2 and A375 #3 due to tight similarity between cell line fluxes. **(E)** PCA of fluxes from standard FBA for cell line models grown in media conditions of DMEM, MEM, and RPMI. Equivalent to Fig. 2B with HPLM media condition removed prior to analysis.

**Fig. S3.**
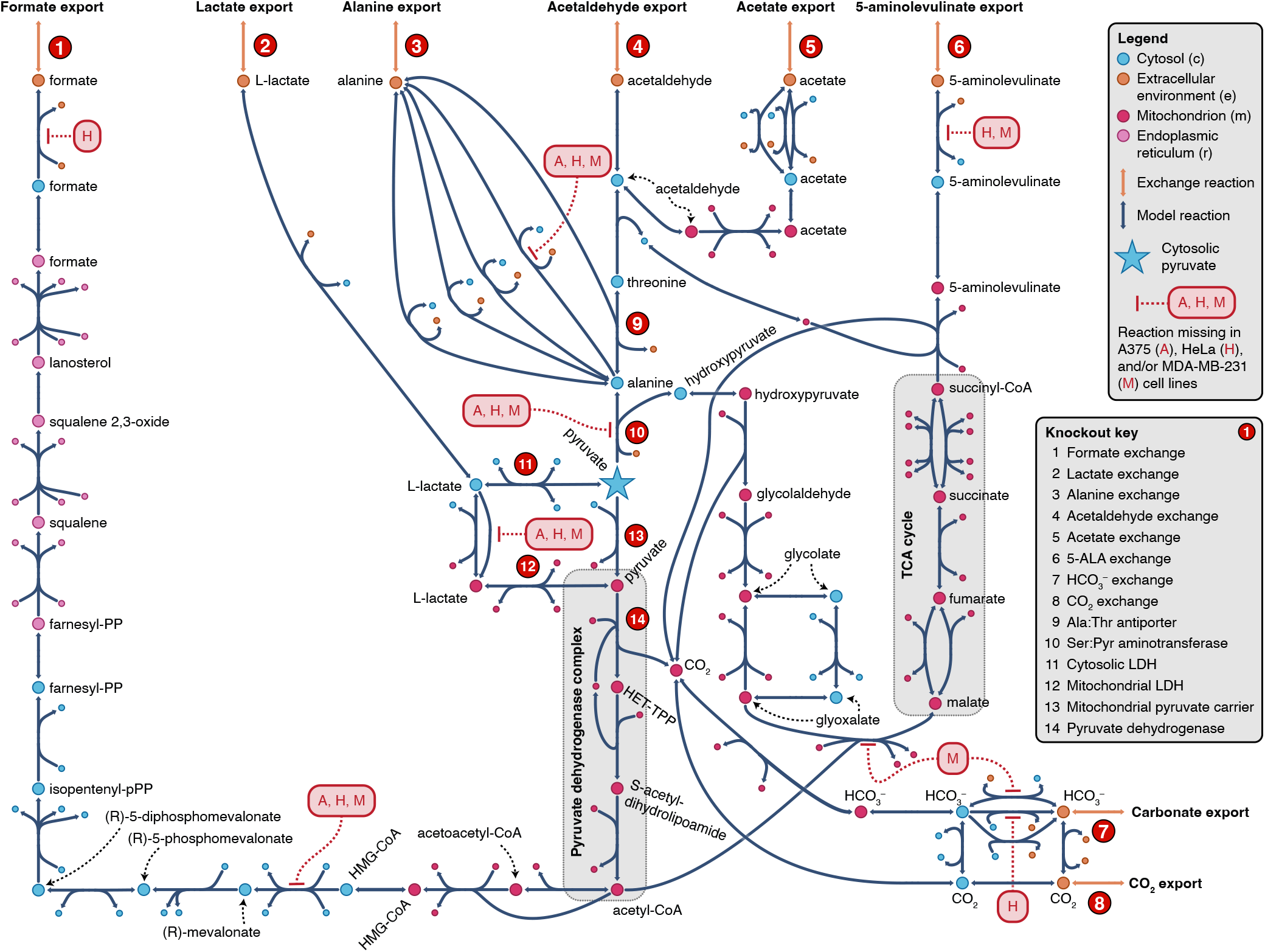
Metabolic map of downstream metabolites generated from pyruvate in HumanGEM. Escher map of carbon-containing exported metabolites downstream of cytosolic pyruvate (denoted as the blue star in the center of the map). Metabolite point colors denote compartmentalization, while arrow color denotes whether the reaction is a reaction between two metabolites in the model or is an exchange reaction at the boundaries of the model. Reactions targeted in the pyruvate knockout screen are denoted with red circles and labeled in the legend. Reactions missing from a given cell line model are denoted for A375 (A), HeLa (H), and MDA-MB-231 (M) with arrows to the relevant reactions.

**Fig. S4.**
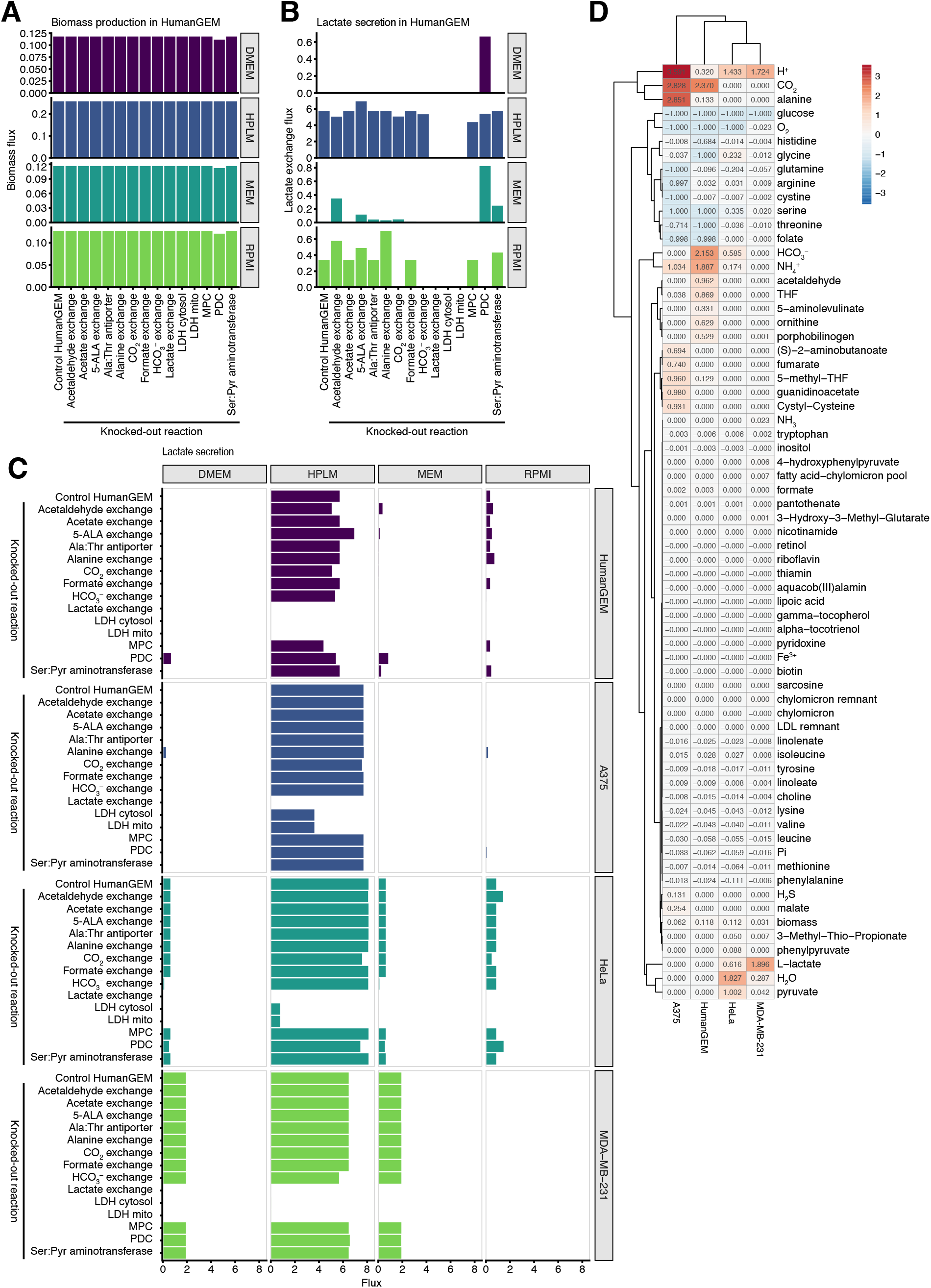
Comparison of HumanGEM and cell line models in DMEM and other media formulations. **(A)** Biomass flux rate in control HumanGEM across media formulations with knockouts of different reactions downstream of cytosolic pyruvate, as shown in Fig. S3. **(B)** Lactate secretion in same FBA conditions as (A). **(C)** Lactate secretion in knockout screen of pyruvate-associated reactions across cell line models and media conditions. **(D)** Clustering heatmap of exchange reactions in DMEM across cell line models. Reactions are denoted by the exchanged metabolite.

**Fig. S5.**
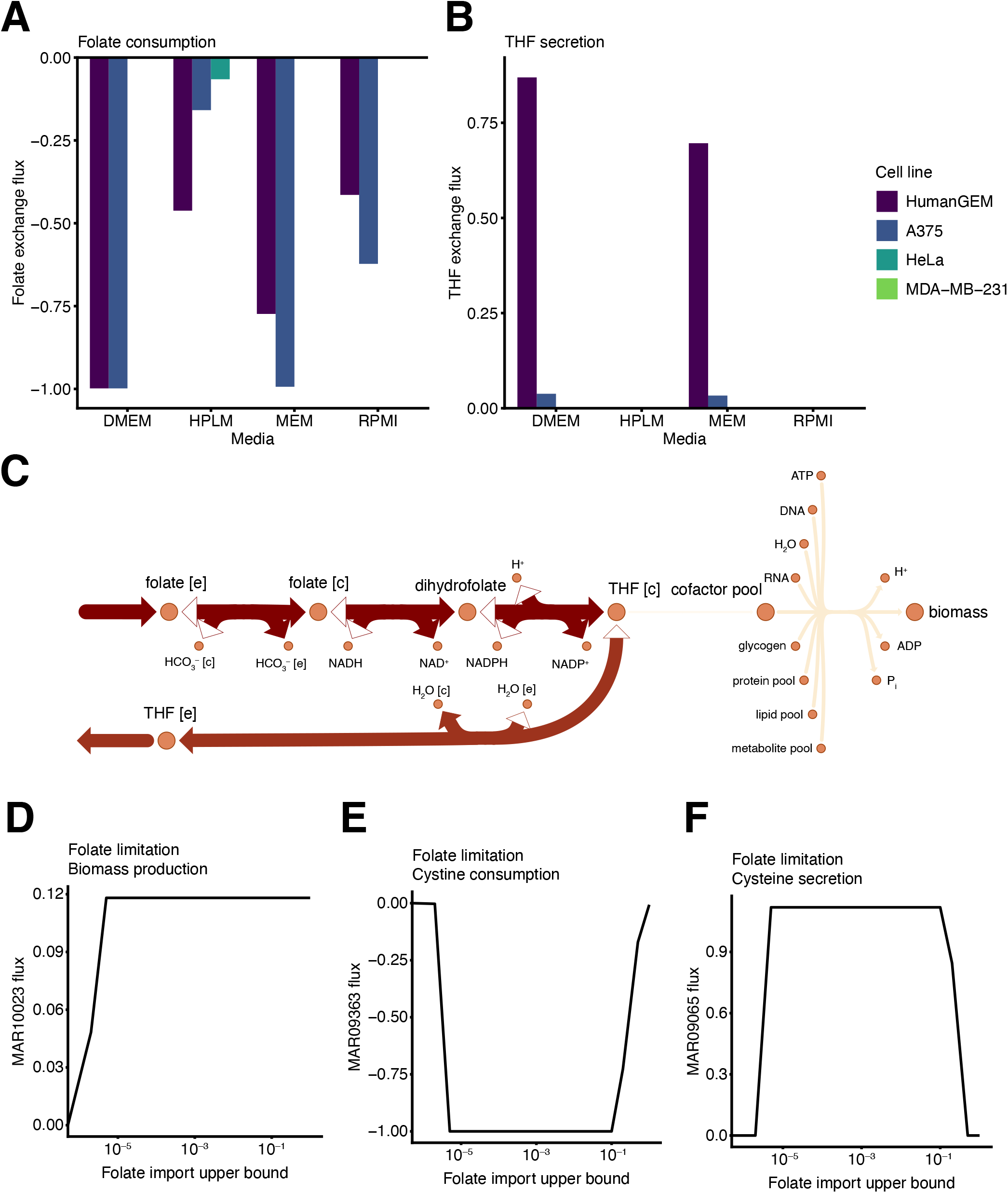
Folate is consumed at *V*_max_ in control HumanGEM to balance redox state. **(A)** Folate exchange flux across cell lines and media conditions. Negative exchange flux denotes consumption of the metabolite by the model. **(B)** Secretion of tetrahydrofolate (THF) across cell lines and media conditions. **(C)** Escher map of folate conversion into THF and subsequent export of THF from the cell. Flux values are denoted by reaction arrow color. **(D)** Biomass flux rate across different values of folate limitation (progressive reduction in upper bound of folate import). **(E-F)** Net exchange of (E) cystine and (F) cysteine under conditions of folate limitation.

**Fig. S6.**
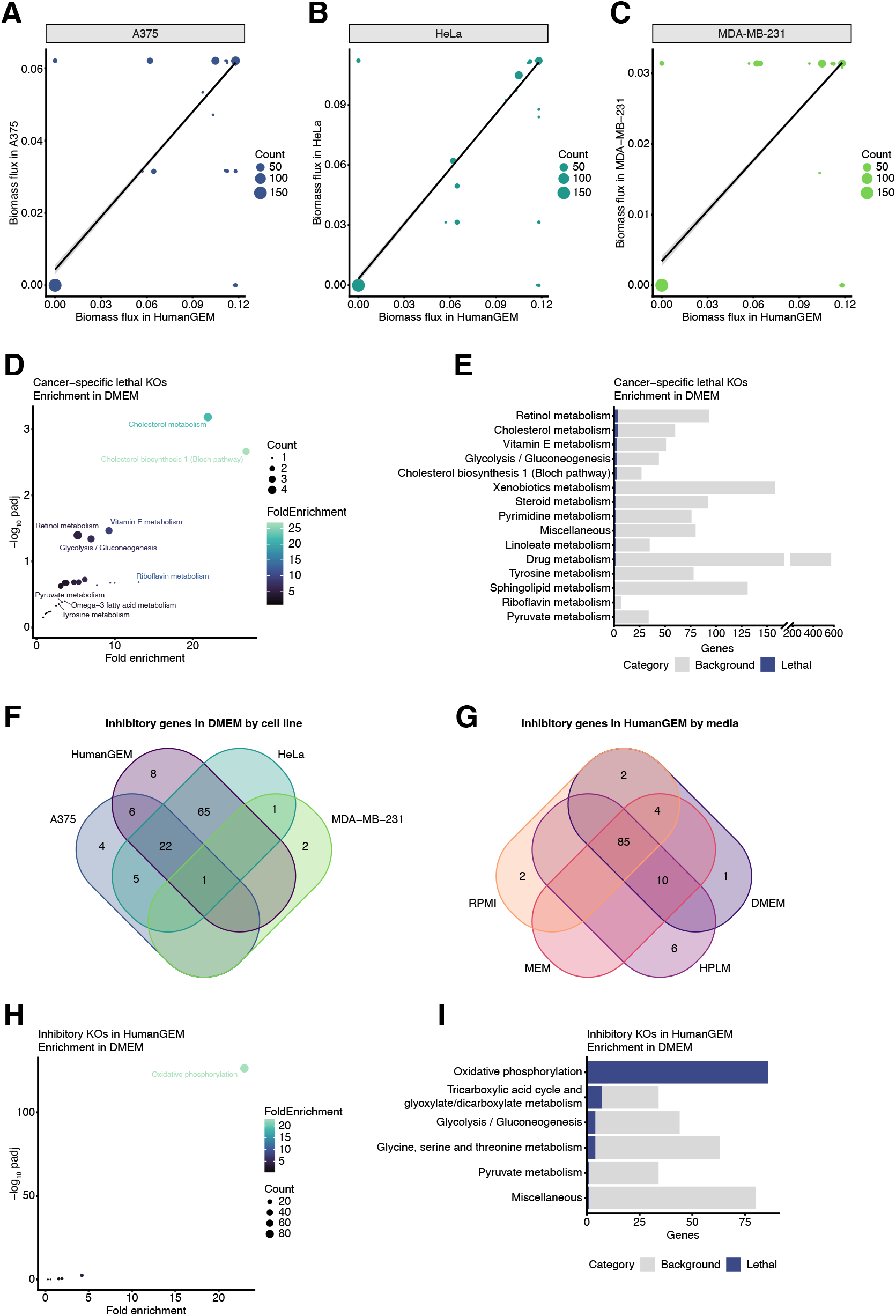
Comparison of gene knockout predictions in HumanGEM and cancer cell lines. **(A–C)** Correlation of biomass flux values for gene knockouts in control HumanGEM and (A) A375, (B) HeLa, and (C) MDA-MB-231. **(D)** Pathway enrichment analysis of lethal knockouts in one or more cancer cell line without lethality in control HumanGEM (refer to Fig. 3E). **(E)** Top pathways from pathway enrichment in (D) by total number of lethal genes in pathway. Total number of genes in pathway is denoted by sum of blue and grey bars. **(F)** Venn diagram of inhibitory genes (where knockout produces a biomass rate greater than 0 but no more than 99% of control biomass) in DMEM by cell line. **(G)** Venn diagram of inhibitory genes in control HumanGEM by media. **H** Pathway enrichment of inhibitory KOs in HumanGEM in DMEM condition. **(I)** Top pathways from pathway enrichment in (H) by total number of lethal genes in pathway. Total number of genes in pathway is denoted by sum of blue and grey bars.

## Notes

### Competing Interest Statement

The authors have declared no competing interest.

